# MARTi: a real-time analysis and visualisation tool for nanopore metagenomics

**DOI:** 10.1101/2025.02.14.638261

**Authors:** Ned Peel, Samuel Martin, Darren Heavens, Douglas W Yu, Matthew D Clark, Richard M Leggett

## Abstract

The emergence of nanopore sequencing technology has the potential to transform metagenomics by offering low-cost, portable, and long-read sequencing capabilities. Furthermore, these platforms enable real-time data generation, which could significantly reduce the time from sample collection to result, a crucial factor for point-of-care diagnostics and bio-surveillance. However, the full potential of real-time metagenomics remains largely unfulfilled due to a lack of accessible, open-source bioinformatic tools.

We present MARTi (Metagenomic Analysis in Real-Time), an innovative open-source software designed for the real-time analysis, visualisation, and exploration of metagenomic data. MARTi supports various classification methods, including BLAST, Centrifuge, and Kraken2, letting users customise parameters and utilise their own databases for taxonomic classification and antimicrobial resistance analysis. With a user-friendly, browser-based graphical interface, MARTi provides dynamic, real-time updates on community composition and AMR gene identification.

MARTi’s architecture and operational flexibility make it suitable for diverse research applications, ranging from in-field analysis to large-scale metagenomic studies. Using both simulated and real-world data, we demonstrate MARTi’s performance in read classification, taxon detection, and relative abundance estimation. By bridging the gap between sequencing and actionable insights, MARTi marks a significant advance in the accessibility and functionality of real-time metagenomic analysis.

## Introduction

Metagenomics is revolutionising our understanding of the diversity and ecology of ecosystems in environmental and even in clinical settings. Advances in this field are largely driven by developments in DNA sequencing technology and the associated analysis tools and pipelines (Escobar-Zepeda et al. 2015). In contrast to the relatively bulky and expensive sequencing platforms developed by Illumina and Pacific Biosciences (PacBio), the MinION, by Oxford Nanopore Technologies (ONT), is a low-cost, portable sequencing device that almost any research group has the resources to acquire and operate. The technology is becoming increasingly used for detecting and characterising pathogenic organisms (Taxt et al. 2020), outbreak surveillance (Quick et al. 2016; Yakovleva et al. 2022), and in-situ sequencing (Burton et al. 2020; Latorre-Pérez et al. 2021), i.e., taking the sequencer to the sample. Significantly, the ONT platform is the first to enable true real-time analysis, with reads able to be accessed during a sequencing run (Leggett and Clark 2017), which combined with the manufacturers API enables focussing of sequencing on specific species within a metagenomic sample using “adaptive sampling” (Martin et al. 2022). Real-time analysis could facilitate a much faster route from sample to results in time-critical situations such as point-of-care diagnostics and bio-surveillance. However, the full potential of real-time sequencing remains largely unrealised due to the lack of open-source, offline, analysis tools.

ONT’s own EPI2ME platform is currently the most well-known example of a real-time metagenomic analysis tool. Initially, ONT developed the EPI2ME Agent software, which featured several cloud-based analysis workflows including two popular metagenomic pipelines: WIMP (What’s In My Pot) and ARMA (Antimicrobial Resistance Mapping Application) (Juul et al. 2015). WIMP classified reads using Centrifuge (Kim et al. 2016) with a database of bacterial, viral, and fungal RefSeq genomes and presented a taxonomic tree view of the sample. ARMA used Minimap2 (Li 2018) to align reads against all reference sequences available in the Comprehensive Antibiotic Resistance Database (CARD), to identify antimicrobial resistance (AMR) genes (Alcock et al. 2020). However, due to EPI2ME Agent’s closed nature, lack of flexibility or customisation, including classification parameters and choice of reference databases, and the need for a fast and stable internet connection, EPI2ME Agent was unsuitable for many in-field sequencing applications. Subsequently, ONT released the EPI2ME Labs tool (now simply called “EPI2ME”), a collection of Nextflow workflows that can be run either locally or in the cloud. This includes wf-metagenomics, a taxonomic classification pipeline that can use either Kraken2 (Wood et al. 2019) or Minimap2 with a user-provided database. Despite these advancements, EPI2ME still has limited data visualisation and data export capabilities, and no between-run comparison features.

Here, we present MARTi, Metagenomic Analysis in Real-Time, an open-source software tool that enables real-time analysis, visualisation, and exploration of metagenomic sequencing data. MARTi allows users to choose a classification method (Kraken2, Centrifuge, BLAST (Camacho et al. 2009), or DIAMOND (Buchfink et al. 2021)), customise classification parameters, and provide their own databases. As an offline tool with low memory options, MARTi can be used for in-field taxonomic composition and AMR gene analysis in real-time on a standard laptop. MARTi can also carry out larger scale, complex analyses using a high performance computing (HPC) cluster. Finally, MARTi features an intuitive browser-based GUI that facilitates exploration of metagenomic results and enables intuitive side-by-side sample comparison on a laptop/desktop, tablet, or smartphone. An installation-free demo of the MARTi GUI is available at https://marti.cyverseuk.org/.

## Results

### Software architecture

MARTi consists of two main components: the MARTi Engine, a Java backend that performs the analysis of the sequencing data; and the MARTi GUI, an easy-to-use browser-based graphical user interface for visualising, exploring, and comparing results (Figure 1). The modularity of MARTi allows it to be configured in different ways depending on the needs of the experiment and the computational equipment available.

**Figure 1.**
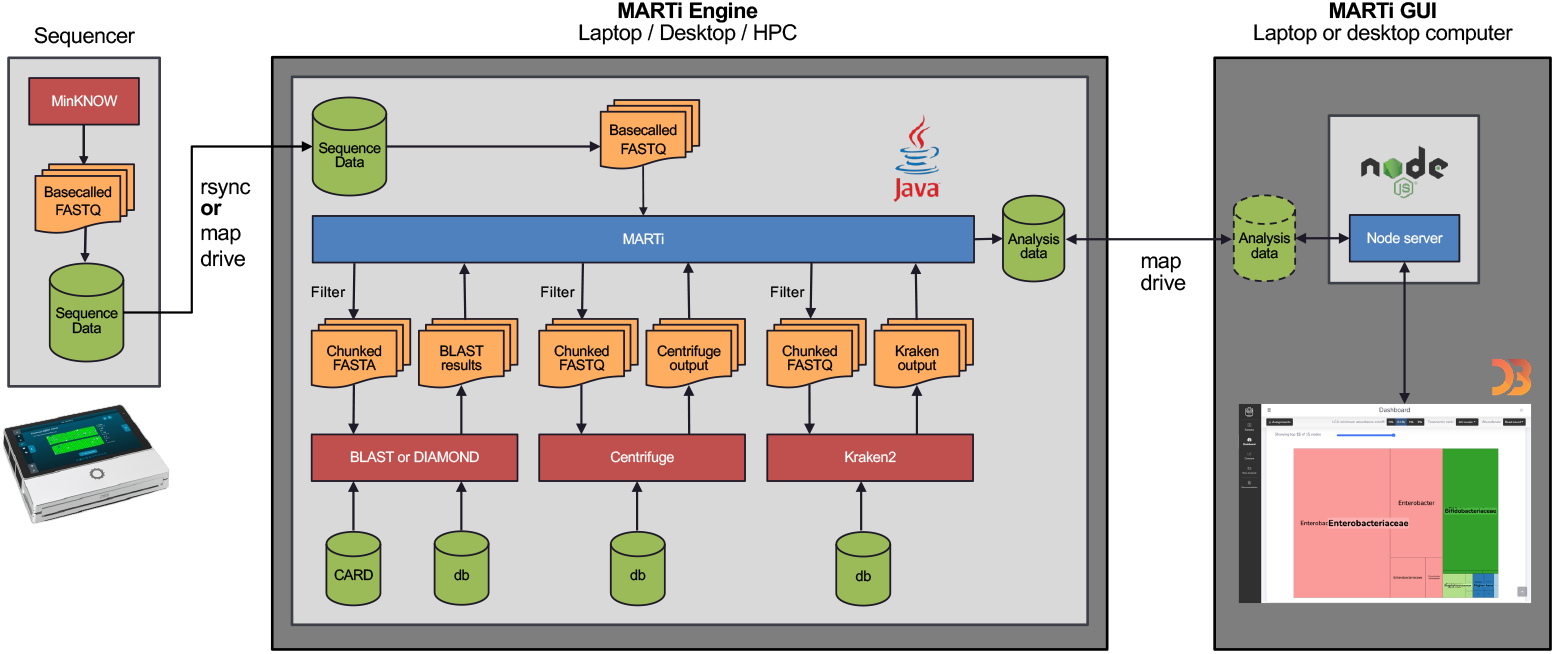
MARTi software architecture. MARTi consists of two main components: the MARTi Engine, a Java backend that performs the analysis of the sequencing data; and the MARTi GUI, a browser-based graphical user interface for visualising, exploring, and comparing results.

In the local configuration, both the Engine and GUI are installed on a single device. This configuration does not rely on any external computing resources and is suitable for in-situ analysis. A nanopore sequencing device, such as a MinION or GridION, generates batches of base-called reads that are accessible to the MARTi computer either by mapping the sequencer’s drive or via the *rsync* utility. The MARTi Engine, initiated through the command line or via the MARTi GUI, analyses new data as it becomes available from the sequencer. The GUI server is run on the same computer and provides analysis results to any connected web browser, which could be a browser on the same computer or any other device, including tablets, on the same network.

In the HPC configuration, the MARTi Engine runs on an HPC or separate server, whilst MARTi GUI resides elsewhere, for example on a laptop/desktop (Fig 1). MARTi supports job scheduling via SLURM (Simple Linux Utility for Resource Management) on the HPC or its own process-based job scheduling (when SLURM is not available). This approach allows greater parallelisation of analysis processes, facilitating queries against larger databases (including NCBI’s nt nucleotide database), and enabling analysis of multiple runs simultaneously. The Engine analyses the data on the HPC or separate server and generates output files for the front end. The MARTi GUI’s server is run on a desktop/laptop that has access to the network location containing the output files.

### MARTi Engine

The MARTi Engine carries out the following main processes: Prefiltering (to remove low-quality and short reads); Classification (assigning reads to taxa); AMR analysis (an optional AMR gene detection step); and Generating (writing output and analysis files for downstream analysis, including those needed by the GUI).

#### Prefiltering

Basecalled reads first pass through a prefilter that removes low-quality or short reads based on user-set thresholds. The default minimum length is 500 bp to remove short reads that have low taxonomic discriminatory power. The default minimum mean quality score is 9, equal to the pass read cutoff of ONT’s High accuracy (HAC) basecalling model. Reads that pass prefiltering are batched into chunks of a specified size for further analysis. The division of reads into chunks permits the parallelisation of the later stages.

#### Classification

The three main ways by which MARTi taxonomically classifies reads are BLAST (Camacho et al. 2009) (followed by its own Lowest Common Ancestor (LCA) algorithm using the NCBI taxonomy - see Methods), Centrifuge (Kim et al. 2016) and Kraken2 (Wood et al. 2019). MegaBlast is the default BLAST algorithm for nucleotide-to-nucleotide comparison within MARTi, but other options are supported via the *blastn -task* option. MARTi also supports translated nucleotide-to-protein comparison (with *blastx* and DIAMOND) and translated nucleotide-to-translated nucleotide (with *tblastx*).

#### AMR analysis

If specified in the configuration file, the MARTi Engine will also BLAST the filtered reads to CARD for AMR gene identification. CARD’s metadata is also used to assign drug class, resistance mechanism, and other information to hits. The host species of an AMR gene hit can sometimes be identified using the taxonomic classification of flanking DNA sequences, a process known as walkout analysis (Leggett et al. 2020). If a read is not long enough to contain significant flanking regions, it is likely to have hits to multiple species and therefore be assigned to a higher taxonomic level by the LCA algorithm. Similarly, AMR genes located on plasmids can pose a challenge as the flanking regions can often have ambiguous taxonomic hits.

#### Generating output

The Engine writes out analysis data and output files, including those required for the MARTi GUI to function. To reduce disk usage, the Engine will compress large intermediate files such as the BLAST output and chunked read files using gzip.

### MARTi GUI

The MARTi GUI is a lightweight browser-based frontend that allows users to view and interact with results generated by the Engine without the need for in-depth bioinformatic knowledge or a strong understanding of the command line (Fig. 2). Crucially, the GUI will update as sequencing progresses and new data becomes available – assuming sufficient compute resources are available, this will appear in near real-time. The GUI has four pages: Samples, for selecting and loading available sample results; Dashboard, for viewing metrics and real-time analysis results of a single sample; Compare, for sample comparison; and New analysis, which allows users to configure and start a local MARTi analysis from the MARTi GUI. All plots on the GUI have options for customisation, update automatically as new data is generated by the Engine, and can be exported as vector (e.g. for papers or presentations) or raster images. Plots on the Dashboard and Compare pages can be displayed at different taxonomic ranks, e.g. family, genus, species, four different LCA minimum abundance cutoffs (0, 0.1, 1, and 2%, where reads from taxa with classified read proportions under the threshold are bumped up the tree until they are in a taxonomic bin that satisfies the minimum), and abundance can be based on read counts or base pair yield.

**Figure 2.**
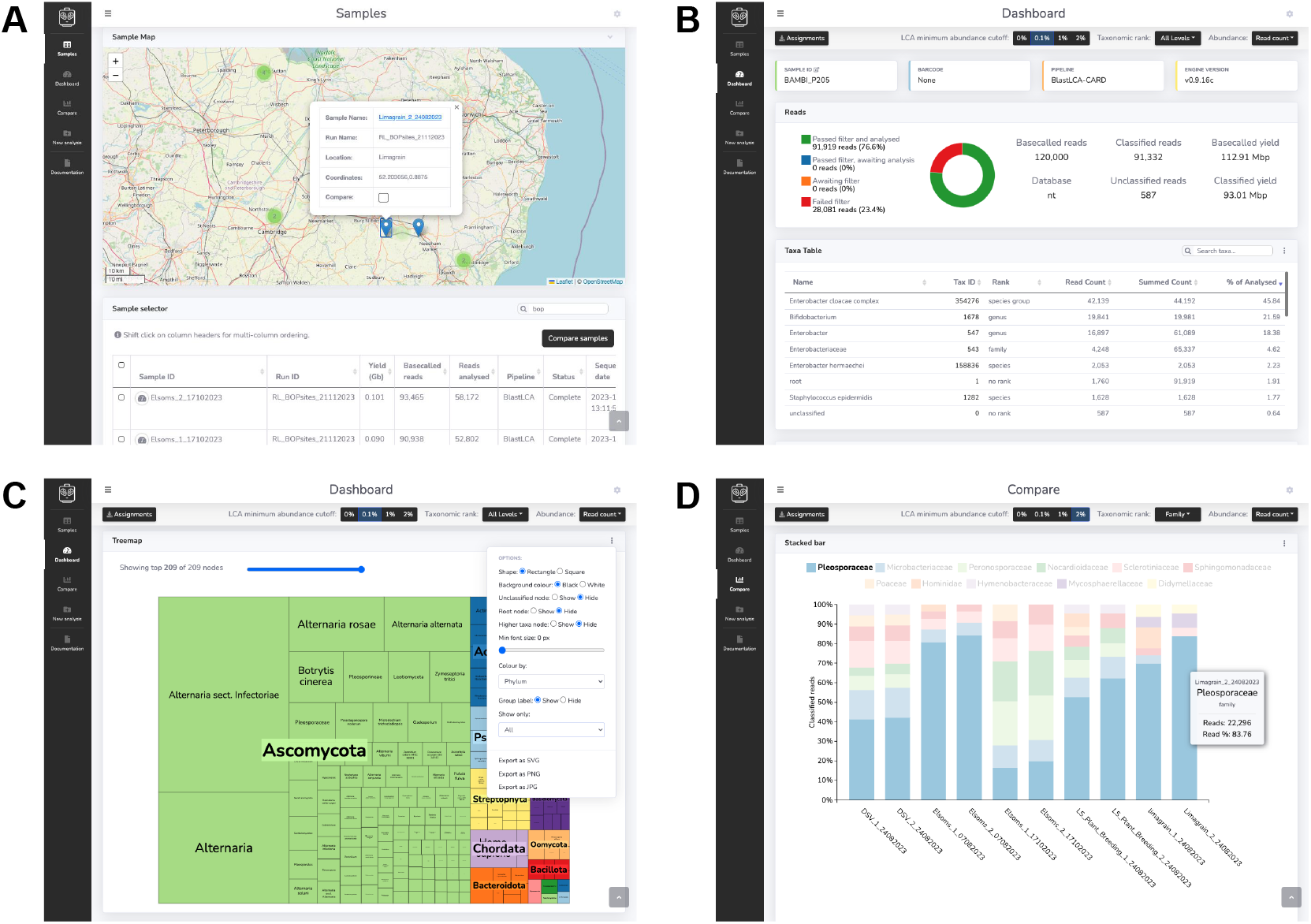
The MARTi GUI. (A) The Samples page is the landing page for the interface and allows users to view and load available samples. (B) The Dashboard is for viewing results of an individual sample. (C) The Dashboard displays analysis results using various tables and interactive plots that update in real time as updates become available. Each plot has an options menu (shown here on the treemap plot) for plot customisation and export buttons. (D) The Compare page can be used to compare taxonomic composition and AMR hits between samples.

#### Samples

The main purpose of the Samples page is to allow users to select and load available samples into Dashboard and Compare analysis modes. However, it also has a data export panel and a sample map (Fig. 2A). The export panel enables users to export selected sample data in the format of their choice for downstream analysis, including formats accepted by other metagenomic tools. The map view plots samples with location metadata on a world map and can be used directly for sample selection. The markers displayed on the map reflect sample table filtering.

#### Dashboard

Taxonomic classifications, AMR hits, and other sample information for an individual sample can be viewed on the Dashboard page (Fig. 2B). Taxonomic results are displayed in a table, and plotted as a donut, taxonomic tree, and treemap (Fig. 2C). The page also features an accumulation curve that shows the number of taxa identified at the selected taxonomic rank over reads analysed. If AMR analysis was carried out the page will also feature a table of AMR hits, a host organism donut plot, and a donut plot of AMR genes, resistance mechanisms, and drug classes associated with species.

#### Compare

This page facilitates comparison of taxonomic composition and AMR presence (Fig 2D). This page features a stacked bar chart, multi-donut plot, combined taxonomic tree, taxonomic heatmap, accumulation plot, and AMR heatmap plot.

#### New analysis

The MARTi Engine requires a configuration file for each analysis, providing all the details for the analysis to be performed. A template configuration can be generated by the MARTi Engine using the *-writeconfig* flag. Alternatively, the New analysis page can be used to initiate local analysis or to generate a configuration file that can be downloaded and used elsewhere. It is not possible to initiate new runs on a remote HPC using the GUI, as varying HPC security and configuration approaches make it impossible to create a one-size-fits-all solution. Therefore, a command line option allows new analyses to be initiated in an HPC environment.

### Taxonomic classification pipeline evaluation

We evaluated the performance of 17 different classification pipelines (Table 1) using an *in silico* mock microbial community consisting of 100k simulated reads with a read length N50 of ∼3.6 kb (Supplemental Table S1). We used three tools to run the pipelines, MARTi, EPI2ME agent, and EPI2ME (formerly EPI2ME Labs). For each of the MARTi pipelines, we compared the effect of setting LCA minimum abundance filters to 0, 0.1, and 1%. Across the experiments, three different read classification methods were deployed, Centrifuge, Kraken2, and MegaBLAST followed by a LCA algorithm.

**Table 1.**
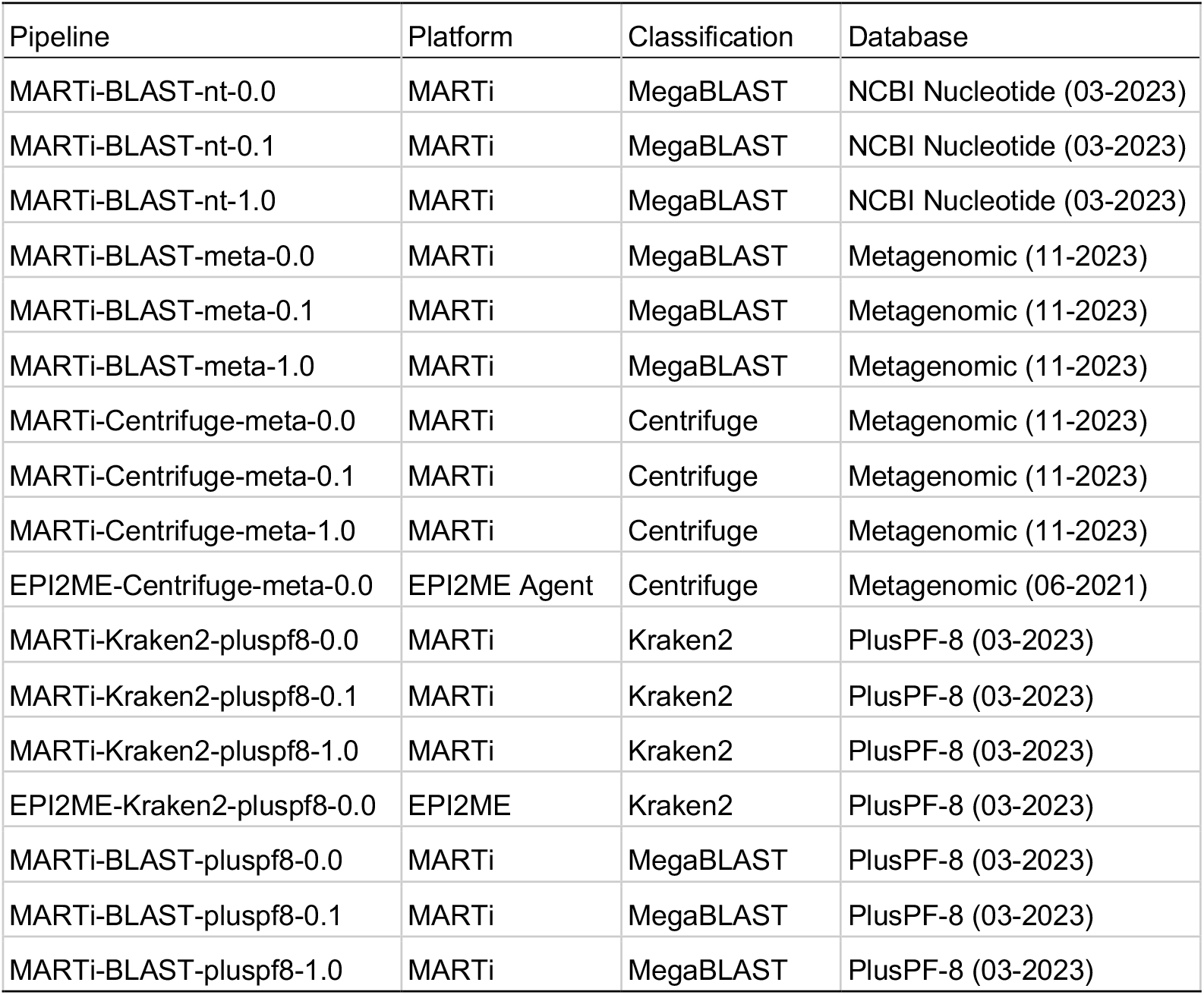
Taxonomic classification pipelines applied to the simulated mock community.

### Read classification

To evaluate the read-level classification performance of each pipeline, we compared the taxonomic classification of each read with its known source (Fig. 3). We assessed classification performance using our simulated metagenomic dataset (proportionally shortened to create a more realistic ∼3.6 kb N50 read length) and the full-length simulated dataset it had been generated from (∼11 kb N50). We also calculated recall, precision, and F-scores, F_1_ and F_0.5_, at genus and species levels (Supplemental Table S2). We define recall as the number of reads correctly classified divided by the total number of reads (100k in this case). Precision was calculated as the number of correctly classified reads divided by the number of assigned reads at a particular taxonomic level and below (i.e., unassigned and reads assigned above a given taxonomic rank are not counted). F-scores were used to summarise the precision and recall scores for each pipeline. F_1_ is the harmonic mean of precision and recall, weighting them equally. F_0.5_ gives more weight to precision, favouring pipelines that minimise false positive classifications.

**Figure 3.**
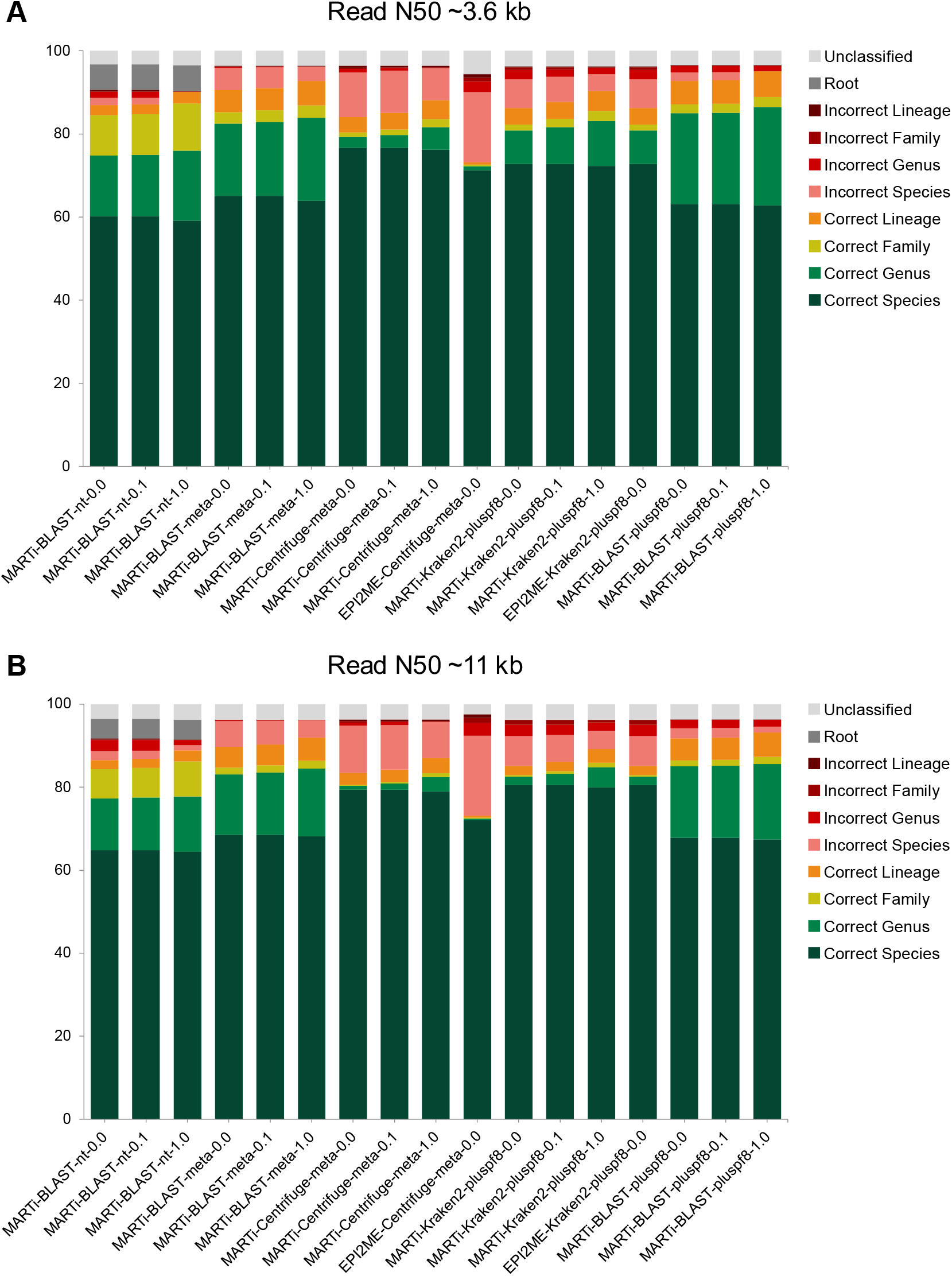
Read classifications for different pipelines using a small mock microbial community consisting of 100k simulated reads. (A) Read classifications using trimmed down reads with a read N50 of ∼3.6kb. (B) Read classification results using the longer reads, read N50 of ∼11kb.

The proportion of reads classified was high across all pipelines, ranging from 94.43% to 96.78% for the shorter readset, and from 96.21% to 97.60% for the longer reads. For both readsets, the EPI2ME-Centrifuge-meta-0.0 pipeline had the highest proportion of incorrectly assigned reads (22.53 to 25.09% for shorter and longer reads respectively), the lowest precision at both species and genus level, and the lowest recall at genus level. Conversely, the MARTi-BLAST-nt-1.0 pipeline had the lowest proportion of incorrectly assigned reads (shorter readset: 0.17%, longer readset: 2.69%), had perfect precision at both species and genus level for the shorter readset and the highest precision at species level for the longer readset. However, the MARTi-BLAST-nt-1.0 method also had the lowest species-level recall for both datasets (shorter readset: 0.59, longer readset: 0.65), and the MARTi-BLAST-nt pipelines were the only methods that classified a considerable proportion of reads to the “Root” node (shorter readset: 6.17%, longer readset: 4.72%).

MARTi-Kraken2-pluspf8-0.0 and EPI2ME-Kraken2-pluspf8 produced identical results, as expected, given that we used the same classification database and version of Kraken2. With both datasets we observed that Kraken2 and Centrifuge-based pipelines had higher species-level recall than BLAST-based pipelines. However, BLAST-based pipelines always had the highest species-level precision for each database used. Regardless of the classification method, implementing a minimum abundance cut-off within MARTi consistently increased the proportion of correctly assigned reads.

### Taxon detection

The ability to accurately detect organisms present in the mock was evaluated across the different pipelines at both species and genus level using the shorter readset (Fig. 4). At each level we calculated recall, precision, and F-scores, F_1_ and F_0.5_ (Supplemental Table S3).

**Figure 4.**
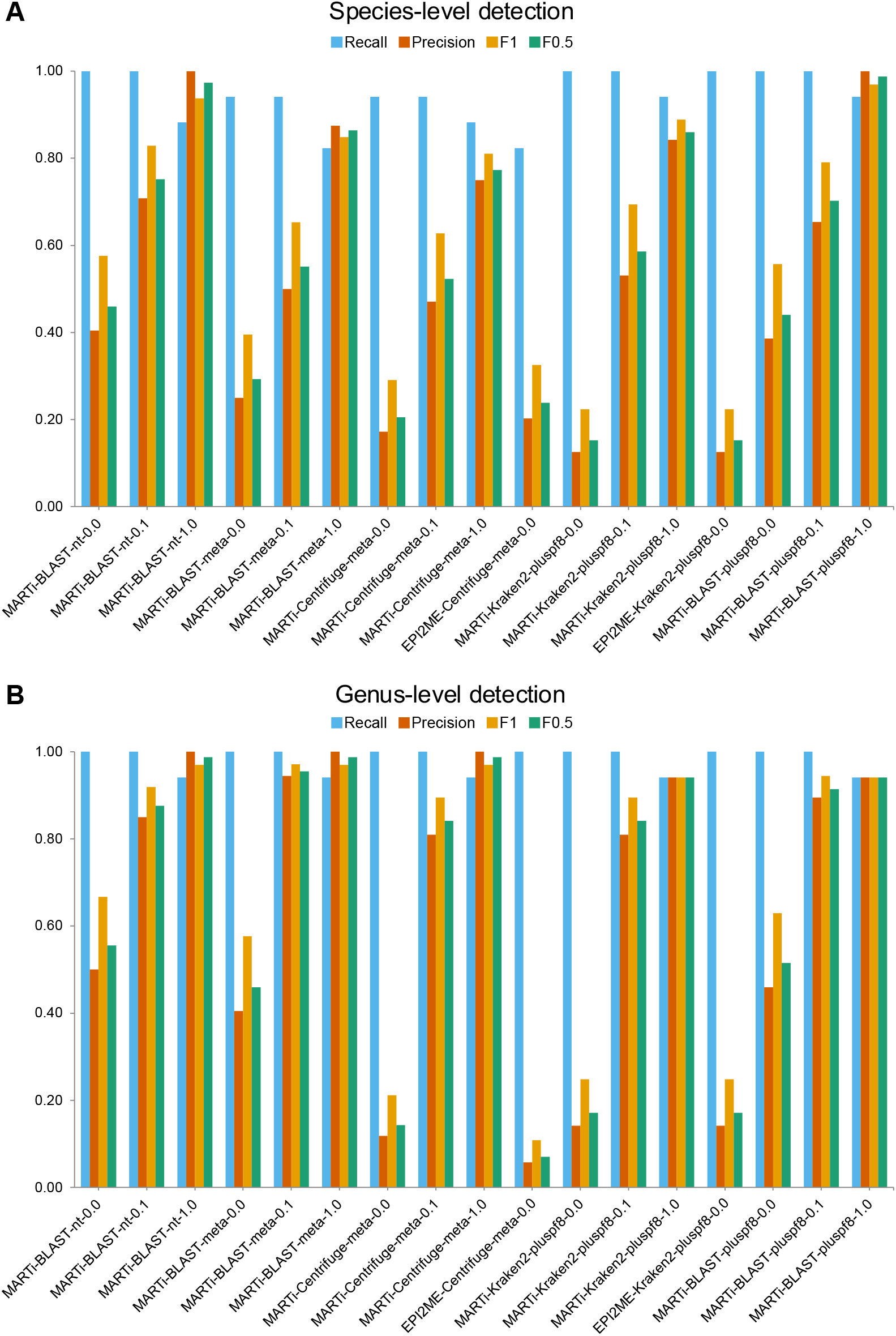
Taxa detection metrics (recall, precision, F1 and F0.5) for different pipelines using simulated reads (Read length N50 ∼3.6 kb) from a small mock microbial community. (A) Species-level detection metrics. (B) Genus-level detection metrics.

Recall was defined as the number of correctly identified taxa divided by the total taxa expected, thus a score of 1 indicates all expected taxa were detected. Precision was calculated as the number of correctly identified taxa at a taxonomic level divided by the total number of taxa identified at that level (true positives + false positives).

At the species level, 7/17 of the methods successfully detected all seventeen species present within the mock community. The MARTi-Blast-meta-1.0 and EPI2ME-Centrifuge-meta-0.0 pipelines had the lowest species-level recall, detecting 14 of the 17 species. MARTi-Centrifuge-meta-0.0 detected a greater number of expected species (16/17) than the EPI2ME-Centrifuge-meta-0.0 pipeline (14/17), likely due to its use of a more recent database. Kraken2-based pipelines without an LCA cutoff had the highest number of false positive species detections. However, the majority of these false positives were supported by very few reads and applying a 1% minimum cut-off removed almost all of them, thereby increasing precision. MARTi-BLAST-nt-1.0 and MARTi-BLAST-pluspf8-1.0 achieved perfect precision scores, with no false positive species detections, and the highest F_0.5_ scores (0.97 and 0.99 respectively).

Genus-level analysis generally yielded higher recall and precision compared to species-level analysis. At this level, 12 out of 17 methods successfully detected all 17 genera in the mock dataset. Similar to species-level results, applying a minimum abundance cutoff at the genus level typically reduced recall while increasing precision. The five pipelines that detected 16 out of 17 genera all used a 1% minimum abundance cutoff and consistently missed the *Clostridium* genus, which had the lowest abundance at just 0.5%. The highest F_1_ and F_0.5_ scores were achieved by MARTi-BLAST-meta-0.1 and MARTi-BLAST-meta-1.0, respectively.

### Relative abundance estimation

We assessed each pipeline’s ability to estimate relative taxonomic abundance based on read counts. The resulting communities were compared using the known composition of the simulated dataset and Bray-Curtis dissimilarities (Fig. 5, Supplemental Table S4). At the species level, abundance estimates by the Kraken2-based methods were the most accurate (based on Bray-Curtis dissimilarity values), followed closely by the MARTi-Centrifuge-meta pipelines. EPI2ME-Centrifuge-meta had the highest dissimilarity and therefore was considered to have the least accurate species-level abundance estimation. The second most relatively abundant species in the simulated dataset, *Faecalibacterium prausnitzii* (14% of the reads), was generally underrepresented by most methods, with values ranging from 0 to approximately 6.7%, with the exception of EPI2ME-Centrifuge-meta, which reported a more accurate relative abundance of ∼14.2%.

**Figure 5.**
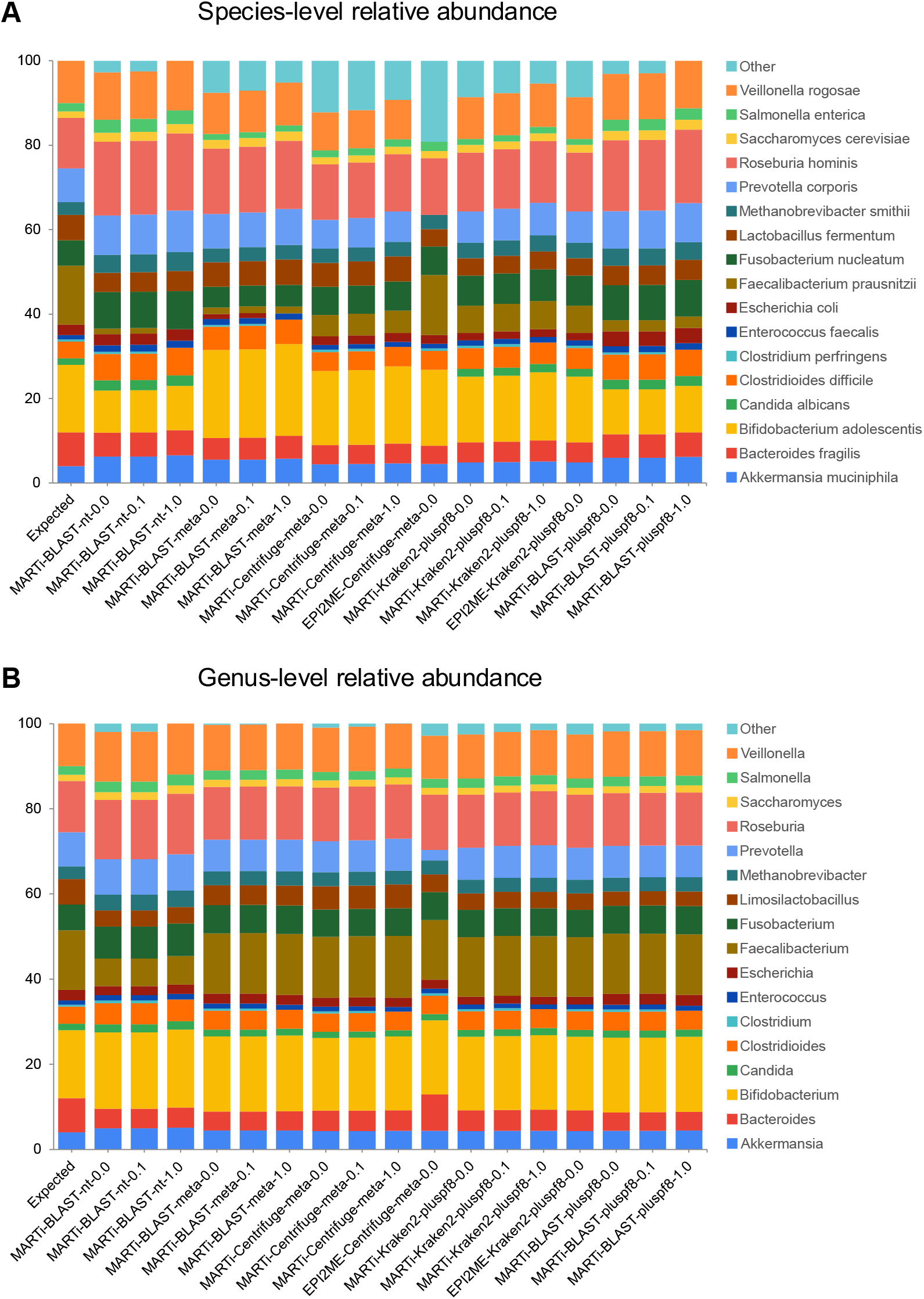
Relative abundance estimates for different classification pipelines using simulated reads from a small mock microbial community at (A) Species-level and (B) Genus-level. The first bar represents the expected abundances of each taxon in the mix based on read counts. False positive classifications are grouped into “Other”.

For every pipeline, relative abundance profiles were more accurate at the genus level than at the species level. At the genus level, MARTi-Centrifuge-meta pipelines had the lowest dissimilarity scores and appeared most like the expected profile. MARTi-BLAST-nt pipelines had the highest dissimilarity scores, most likely due to their underestimation of the *Faecalibacterium* genus (Expected: 14%, MARTi-BLAST-nt pipelines: 6.53-6.70%, All other pipelines: 13.97-14.48%).

### Local configuration analysis rate

We compared the analysis rate of the three main classification methods available within MARTi by analysing 100k simulated mock microbial reads (∼1 Gbp of data) locally on a MacBook Pro (Table 2). In the MARTi configuration files for each method, we set the maximum number of parallel MARTi jobs and specified the number of threads each classification tool could use to optimize the tools’ use of computational resources. Among the methods, Kraken2 was the fastest with a total wall-clock analysis time of 204 seconds, classifying around 29,412 reads per minute (rpm). This was closely followed by Centrifuge, which took 224 seconds, classifying at 26,786 rpm. BLAST was by far the slowest, with a total analysis time of 49,895 seconds (i.e. ∼14hrs or overnight), analysing at a rate of 120 rpm.

**Table 2.** Analysis rate and memory use for the main classification pipelines within MARTi running on a MacBook Pro.

| Pipeline | Jobs | Threads | Peak Memory (MB) | Execution time (s) | Reads per min (RPM) |
| --- | --- | --- | --- | --- | --- |
| MARTi-BLAST-meta | 8 | 1 | 65526 | 47457 | 126.43 |
| MARTi-Centrifuge-prok | 2 | 8 | 41958 | 224 | 26785.71 |
| MARTi-Kraken2-pluspf8 | 2 | 8 | 48142 | 204 | 29411.76 |

### Profiling of preterm infant microbiota

To demonstrate the application of MARTi on real data, we reanalysed four published preterm infant microbiomes (Leggett et al. 2020) using the MARTi-BLAST-nt pipeline. Two of the samples were from healthy individuals (P106 and P116), and the other two from preterms clinically diagnosed with necrotising enterocolitis (NEC) (P205 and P8). We then used MARTi GUI to explore the results (Fig. 6). In accordance with the original study, we found the microbiota of healthy samples was dominated by *Bifidobacteriaceae*, whereas that of NEC patients was dominated by *Enterobacteriaceae* (Fig. 6A). The most abundant taxa were consistent and displayed similar relative abundances to those in the original study. In sample P205 (NEC), the dominant taxon was the *Enterobacter cloacae* complex, and in sample P8 (NEC), it was *Klebsiella pneumoniae* (Fig. 6C).

**Figure 6.**
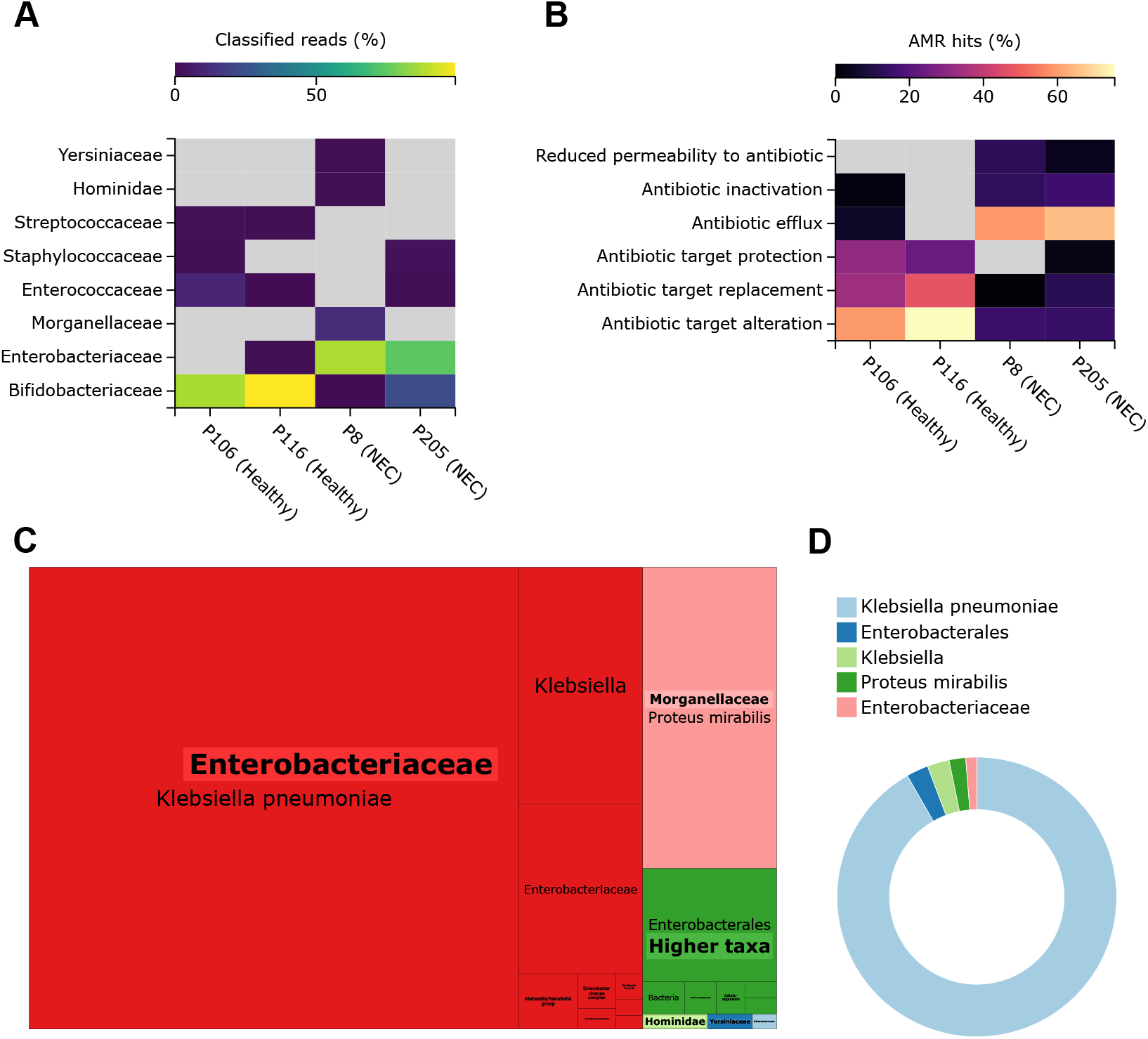
Selected MARTi GUI plots from the reanalysis of four published preterm infant microbiomes. Two of the samples were from healthy individuals (P106 and P116) and the other two from preterms clinically diagnosed with NEC (P205 and P8). (A) Family-level taxonomic classification heatmap. (B) Resistance mechanism composition of the AMR hits within each sample. (C) Treemap of classified reads for P8 (an NEC sample) showing all taxonomic levels grouped by family. (D) Donut plot displaying the proportions of AMR hits for P8 that have been associated with taxa by “walkout” analysis. A 0.1% minimum LCA abundance threshold was used for all the taxonomic classification plots. Plots were exported from the MARTi GUI as SVGs and arranged with Inkscape.

We also ran MARTi’s AMR analysis on the microbiome datasets. We observed distinct differences in AMR resistance mechanism profiles between healthy and NEC samples, with a significant portion of the AMR hits in NEC individuals associated with antibiotic efflux and inactivation mechanisms that were almost entirely absent from the healthy samples (Fig. 6B). Most of the AMR hits within sample P205 (NEC) were associated with *E. cloacae*, whereas within sample P8 (NEC), they were assigned to *K. pneumoniae* (Fig. 6D). We used a more recent version of the CARD database for our reanalysis with MARTi, and as a result, we identified many antimicrobial resistance ontologies (AROs) that were not present in the database used for the original analysis. Notably, 15 of the 32 AROs identified by MARTi in P8 (NEC), including the top four AROs by read count, were absent from the database used in the original analysis.

## Discussion

ONT’s long-read sequencing platforms are the first to enable progressive real-time data analysis, with the potential to revolutionise metagenomics by improving classification accuracy and assembly quality, reducing time to results, and enabling new ways of working, such as bringing the lab to the sample. However, the full potential of nanopore sequencing remains largely unrealised due to the lack of open source, offline, real-time analysis tools and pipelines. In this manuscript, we present MARTi, an open-source software tool that enables real-time analysis and visualisation of metagenomic sequencing data. MARTi provides a rapid, lightweight web interface that allows users to view community composition and identify antimicrobial resistance (AMR) genes in real-time.

MARTi allows users to classify their reads in three main ways: BLAST, followed by a LCA algorithm; Centrifuge; and Kraken2. We have demonstrated and compared each of these methods using simulated gut microbiome reads, where the true origin of each read is known. We found that *k*-mer-based tools, Kraken2 and Centrifuge, had the highest species-level recall. This means that they correctly classified the greatest number of reads to the species level, beating BLAST-based pipelines using equivalent databases. However, the *k*-mer-based methods also incorrectly classified the greatest number of reads at species level, leading to lower precision than BLAST-based methods. The ideal classification pipeline would achieve both high precision and recall, but in practice, classification tools often exhibit a trade-off between these metrics (Portik et al. 2022). For many metagenomic applications, high precision at the expense of recall, as seen with BLAST-based methods, will be more desirable in order to minimise false-positive classifications.

Read classification recall was improved for every method when using the longer simulated reads, ∼11 kb read length N50, with the biggest increase seen in Kraken2-based pipelines at the species level. However, precision was usually slightly reduced using longer reads, with the most notable exception being the Kraken2 pipelines, which saw increases in species-level precision. Similar results have been reported by Pearman et al. (2020), who showed that BLAST and Kraken2 recall was improved by the use of longer reads, but precision only improved for Kraken2.

Within the MARTi GUI, users can view taxonomic classification results with different minimum abundance cut-off values (0, 0.1, 1, and 2%). We demonstrated that applying a cut-off value can improve these classification tools’ overall taxonomic detection results, especially the *k*-mer-based tools, Kraken2 and Centrifuge, that had higher numbers of false positives with no filtering. In concurrence with Portik et al. (2022), we observed that the application of a minimum abundance threshold can dramatically reduce the number of false positive detections, increasing precision, but at the cost of increased false negatives, reducing recall, especially at more stringent threshold values.

We estimated relative taxonomic abundances for each of the analysis pipelines applied to the simulated mock gut microbiome. Abundance was estimated using cumulative read counts, the sum of the reads assigned to a taxon and below, and then we calculated Bray-Curtis dissimilarities to compare each of the estimated mock community compositions to the known composition. Kraken2-based methods produced species-level abundance profiles most similar to the expected profile, while MARTi-Centrifuge-meta pipelines exhibited the lowest dissimilarity scores at the genus level.

A potential limitation of using long-read count data for abundance estimation is that read length variation is not considered. For real metagenomic samples, average read lengths obtained for different species within a mix can vary greatly and make read-based abundance estimates less accurate. To account for this, we have added an option within the MARTi GUI for users to switch between read-count-based abundance and abundance based on cumulative assigned base pairs.

For real-time metagenomic applications, analysis rate should be greater than or equal to sequencing rate to keep up. We demonstrated that MARTi can analyse ∼1 Gbp every 3.5 minutes when using Kraken2 for classification on a MacBook Pro. At this rate, MARTi can analyse the data from a 72-hour MinION run (producing 48 Gbp theoretical max output) in under 3 hours on a laptop. The analysis rates we observed for Kraken2 and Centrifuge were similar to those reported by Marić et al. (2024).

When it comes to sequence comparison, BLAST is often considered the gold standard, but it is also orders of magnitude slower than newer *k*-mer-based tools. We recommend users run MARTi on an HPC or powerful server if they want to use BLAST-based classification. When MARTi classifies reads with BLAST followed by an LCA algorithm, it produces similar classification results and has the same taxonomic resolution as other tools based on this approach.

We re-analysed published infant microbiome datasets with MARTi using MegaBlast against the nt database for classification. The main taxonomic findings were the same, with some differences in the less abundant taxa. This is not surprising considering a similar BLAST-based approach was taken in the original study, with differences in results largely due to growth in the nt database and how MARTi’s LCA algorithm interpreted the BLAST output. We also found that up to ∼40% of AMR genes identified in the re-analysed samples were not previously found due to being absent from the database used in the original study. These results highlight the impact of parameter and database choice on metagenomic classification. One feature of MARTi that is not present in EPI2ME is the sample comparison mode, making it easy to compare the taxonomic assignments and AMR hits across samples. This has obvious uses for comparing biological communities between different sites, conditions and time points. However, the compare mode is also useful for assessing the effect of database and parameter choice on the same dataset.

MARTi represents a significant advance in real-time metagenomics, addressing the need for a flexible, open-source, and offline analysis tool. Through comprehensive evaluation using simulated and real-world datasets, MARTi has demonstrated robust performance in read classification, taxon detection, and relative abundance estimation, leveraging classification tools BLAST, Centrifuge, and Kraken2. Although initially developed for long-read metagenomics, MARTi can be applied to short reads and metabarcoding projects with some parameter changes. The integration of an intuitive, browser-based GUI facilitates accessibility and usability, making metagenomic analysis more attainable for diverse research settings. Furthermore, MARTi’s capability to operate in different configurations ensures scalability and adaptability to varying computational resources, making it a useful tool for real-time in-field metagenomic studies and larger scale post-run analysis. Future enhancements and community contributions will likely expand MARTi’s functionality, further solidifying its role in metagenomic data analysis.

## Methods

### Lowest Common Ancestor Algorithm

When classifying reads using BLAST (nucleotide or protein) or DIAMOND (a BLAST-like protein alignment tool), MARTi implements a Lowest Common Ancestor (LCA) algorithm (see below) to assign reads to taxa based on the alignment results. This algorithm assigns reads to the lowest taxonomic level consistent with “good” hits. The definition of good is configurable and depends on the BLAST bit score, length of match, percent identity of the match and the maximum number of hits to consider. MARTi implements a Lowest Common Ancestor algorithm as follows:

1. Reads are BLASTed against a user-defined database. This may be, for example, the whole of NCBI nt, a bacteria subset, RefSeq genomes, proteins such as NCBI’s nr, or a custom database. This results in a set of between 0 and many hits for each read.
2. For a given read, the set of “good hits” is identified by finding the highest scoring hit (according to the BLAST bitscore), then finding all hits with a score within 90% (default value, but configurable) up to an optional limit (default 100 hits, but configurable). Note, if the number of hits considered is limited, there is a risk that the first X hits may not contain the best hit; the larger this number, the lower the risk of misclassification.
3. For each good hit, the taxonomic path is determined by referring to the NCBI taxonomy. For example:

~~~
“root, cellular organisms, Bacteria, Proteobacteria,
Gammaproteobacteria, Enterobacterales, Enterobacteriaceae,
Klebsiella, Klebsiella pneumoniae”
~~~

The taxonomic paths for all good hits are compared to determine the common ancestor. This involves starting at the root node and working downwards, comparing nodes (first “root”, then “cellular organisms”, then “bacteria” etc.) until paths diverge. The last node in common before paths diverge is the lowest common ancestor and the read is assigned to this taxon.

### Mock microbial community read simulation

We generated a small mock microbial community dataset consisting of 100k simulated nanopore reads to evaluate the classification pipelines available within MARTi (Supplemental Table S1). Long reads were simulated from 17 RefSeq genomes (15 prokaryotes and 2 eukaryotes) using NanoSim v3.1.0 (Yang et al. 2017), subsampled, then combined and randomly shuffled. The original simulated data set has a total of 1,041,760,674 base pairs (2.09 GB of FASTQ files) and an N50 read length of 10,952 bp. To generate a more realistic metagenomic dataset we reduced each read to one third of its original length, resulting in a read length N50 of 3,614 bp.

### Reference databases

We chose commonly used databases for each of the classification tools available through MARTi. For BLAST, we used NCBI’s nucleotide database (nt), a very large and comprehensive collection of nucleotide sequences. The nt database was downloaded as a prebuilt BLAST database from the NCBI FTP site (https://ftp.ncbi.nlm.nih.gov/blast/db). We used the March 2023 version for taxonomic classification pipeline evaluation and March 2024 version for preterm infant microbiota analysis.

EPI2ME agent’s WIMP workflow (v2023.06.13-1865548) uses Centrifuge to classify reads to a metagenomic database consisting of archaea, bacteria, viral, fungal, and human RefSeq genomes (downloaded June 2021). For the MARTi-Centrifuge-meta pipeline, we downloaded a more recent version of this database (Nov 2023) using the following commands within Centrifuge v1.0.3:

~~~
centrifuge-download -o taxonomy taxonomy
centrifuge-download -o library -m -d “archaea,bacteria,viral,fungi”
refseq > seqid2taxid.map
~~~

The human genome was added as follows:

~~~
centrifuge-download -o library -d “vertebrate_mammalian” -a
“Chromosome” -t 9606 -c ‘reference genome’ refseq >> seqid2taxid.map
~~~

The sequences were combined:

~~~
find library -name ‘*.fna’ -exec cat {} >> input-sequences.fna \\;
~~~

Then the centrifuge index was built with the following command:

~~~
centrifuge-build -p 100 --conversion-table seqid2taxid.map --
taxonomy-tree taxonomy/nodes.dmp --name-table taxonomy/names.dmp
input-sequences.fna metagenome
~~~

We built a BLAST database from the same sequences as the Centrifuge database for the MARTi-BLAST-meta pipeline for a more direct comparison of classification performance between Centrifuge and BLAST using the following command:

~~~
makeblastdb -in input-sequences.fna -parse_seqids -blastdb_version 5
-title “refseq_metagenomics” -dbtype nucl -taxid_map taxid_map.txt
~~~

The EPI2ME-Kraken2 pipeline classified the simulated reads with the wf-metagenomics workflow (v2.8.0). The database used for all Kraken2 pipelines was PlusPF-8 (March 2023, downloaded from https://benlangmead.github.io/aws-indexes/k2), a pre-built Kraken2 index containing references for archaea, bacteria, viral, plasmid, human, UniVec_Core, protozoa and fungi. A BLAST database was built using the same sequences for the MARTi-Kraken2-pluspf8 pipeline with the following command:

~~~
makeblastdb -in sequences.fasta -parse_seqids -blastdb_version 5 -
title “k2_pluspf_08gb” -dbtype nucl -taxid_map taxid_map.txt
~~~

### Pipeline read classification

We calculated the recall and precision at genus and species levels for each of the classification pipelines. We define recall as the number of reads correctly classified divided by the total number of reads (100k in this case). Precision was calculated as the number of correctly classified reads divided by the number of assigned reads at a particular taxonomic level and below (i.e., unassigned and reads assigned above a given taxonomic rank are not counted). We also calculated F-scores as a way of summarising the recall and precision information. The F_1_ score is the harmonic mean of precision and recall, providing an equally weighted view of the recall and precision. When calculating the F_0.5_ score, we emphasise precision, placing more importance on minimising false positive classifications. The highest value for either F-score is 1, indicating perfect precision and recall.

### Taxa detection

We scored the presence/absence of taxa for each classification pipeline. We calculated recall, precision, and F-scores for taxa detection at genus and species levels. In this context, we define recall as the number of identified mock taxa divided by the expected number of taxa. Precision was calculated as the number of mock taxa identified divided by the total number of taxa detected at the species or genus level.

### Relative abundance estimation

Relative abundances were estimated for each of the classification pipelines at the species and genus level. For all pipelines, we estimated abundance using cumulative read counts for each taxon, which is the sum of the reads assigned to that taxon and below. The sum of the cumulative counts for false positive taxa were grouped as “Other”. The counts were then normalised as percentages. Bray-Curtis dissimilarity was calculated to compare each of the estimated mock community compositions to the known composition.

### Local configuration analysis rate

We analysed the simulated mock microbial community data with MARTi running in local configuration on a MacBook Pro (2019, 8-core Intel Core i9-9880H @ 2.3 GHz, 64 GB DDR4 RAM). The reads were classified with each of the three main methods available in MARTi: BLAST, Centrifuge, and Kraken2. For BLAST, we used a metagenomic database downloaded with the centrifuge-download command (described earlier in Reference databases). A prebuilt index containing RefSeq prokaryotes was used for Centrifuge classification (April 2018, downloaded from https://benlangmead.github.io/aws-indexes/centrifuge). For the Kraken2 pipeline, we used the pre-built PlusPF-8 index described previously.

### Profiling of preterm infant microbiota

To demonstrate MARTi on real data, we re-analysed four published preterm infant microbiome datasets (Leggett et al. 2020), two from healthy individuals (P106 and P116), and two from preterms clinically diagnosed with NEC (P205 and P8). As described in the original paper, the sequence data is available under ENA accession PRJEB22207. We analysed the first 120k basecalled reads of P106, P116, and P205, and the first 100k reads of P8 using MARTi v0.9.16. Reads passing MARTi’s prefilter, minimum length 500 bp and minimum read quality score of 9, were taxonomically classified using the BLAST pipeline with NCBI’s nt database (March 2024). Additionally, AMR gene analysis was carried out using the CARD v3.2.7 database. Taxonomic assignments and AMR analysis results were explored with the MARTi GUI.

### Data access

The latest version of the MARTi software is available from https://github.com/richardmleggett/MARTi under the MIT License. Documentation can be found at https://marti.readthedocs.io/en/latest/. Additionally, an installation-free demo of the MARTi GUI is available at https://marti.cyverseuk.org/.

The simulated reads used in this study, including both the full-length and one-third length datasets, have been deposited in Zenodo and can be accessed at https://doi.org/10.5281/zenodo.14260487.

## Acknowledgements

The authors acknowledge the support of the Biotechnology and Biological Sciences Research Council (BBSRC), part of UK Research and Innovation; this research was funded by TRDF award BB/R022445/1; Core Capability Grant BB/CCG1720/1; Core Strategic Programme Grant (Genomes to Food Security) BB/CSP1720/1 and its constituent work package BBS/E/T/000PR9817; Strategic Programme Grant (Decoding Biodiversity) BBX011089/1 and its constituent work package BBS/E/ER/230002A. NP was also supported by the BBSRC Norwich Research Park Biosciences Doctoral Training Partnership (grant number BB/M011216/1).

Part of this work was work delivered via the Scientific Computing group, as well as support for the physical HPC infrastructure and data centre delivered via the NBI Research Computing (RC) group.

The installation-free demo of the MARTi GUI was supported by CyVerse UK, a service provided by the Earlham Institute (EI) National Capability in e-Infrastructure (NC3), funded by the BBSRC (BBS/E/T/000PR9814, BB/R000662/1, BB/M018431/1).

## Author Contributions

RML conceived the software. RML, MDC, and DWY supervised NP’s PhD during which the software was initially developed. RML, SM, and NP developed the software. NP and DH performed the experiments. NP analysed the data. NP and RML prepared the manuscript. All authors read and approved the final manuscript.

